# MS2extract: an R Package for a Scalable LC-MS/MS Library Creation

**DOI:** 10.1101/2025.10.05.680545

**Authors:** Cristian Daniel Quiroz-Moreno, Jessica L. Cooperstone

**Affiliations:** Horticulture and Crop Science Department, The Ohio State University, Columbus, USA, 43210; Food Science and Technology Department, The Ohio State University, Columbus, USA, 43210

## Abstract

Reference MS/MS libraries are useful to aid in metabolomics compound identification. However, creating these libraries can be challenging because of the complexity of MS/MS data handling, the need for manual spectrum inspection, and requirement for custom-designed bioinformatics tools for this purpose. Although licensed and open-source alternatives to create MS/MS reference libraries exist, most of these pipelines require manual extraction of each reference spectrum, which is a time-consuming and error-prone process. Here, we present MS2extract, an R package facilitating the user-led creation and automation of MS/MS spectral libraries. The core MS2extract workflow consists of importing MS/MS raw data, detecting specific precursor ions, extracting MS/MS spectra, removing low-intensity signals, and exporting a spectral database. In addition, libraries created with MS2extract are compatible with GNPS2 batch upload to maximize MS/MS library reuse. We used MS2extract to create PhenolicsDB, a phenolics-focused MS/MS library with fragmentation patterns of 71 authentic analytical standards collected in positive and negative polarity, and multiple collision energies, with 320 MS/MS reference spectra. Finally, we employed PhenolicsDB to identify apple fruit phenolics. MS2extract and PhenolicsDB are free and publicly available for download and use, and PhenolicsDB is also available in GNPS2.

## Introduction

Tandem mass spectrometry (MS/MS) is a powerful technique to aid in small molecule structural characterization. MS/MS provides information about product ions that result from the fragmentation of a selected ion, which can be distinctive and aid in the identification of chemical species. If MS/MS spectra are collected for known compounds, the spectrum can serve as a reference for future compound identification. A collection of reference MS/MS spectra can be compiled to create libraries, which can later be used for reference to unknown spectra. Significant efforts have been made to create publicly available and licensed MS/MS libraries to aid compound identification, including MassBank^1^, MONA^2^, GNPS libraries^3^, HMDB^4^, Metlin^5^ and mzCloud (<u>Thermo Fisher Scientific</u>). Some public repositories (e.g., GNPS, MONA, and MassBank) support MS/MS library submissions from users which allows the community to benefit from efforts made to compile reference spectra.

Although there are open-source software for mass spectrometry data processing and analysis (e.g., openMS^6^, matchms^7^, xcms^8^, and mzR^9^), only some have dedicated modules to create in-house libraries (e.g., MetID^10^ and MZmine^11^). However, none allow users to perform all of the following: build in-house MS/MS libraries from multiple spectra at once, and export these libraries in an open-source format for other researchers to use that are compatible with MS/MS repository submission. As a result, users must create their own in-house pipelines or manually process MS/MS data, making this a time-intensive and error-prone task.

Here, we present MS2extract, an open-source R package that automates building in-house MS/MS small molecule libraries from raw data. MS2extract offers the option to export libraries in .msp and .mgf formats, which are compatible with most mass spectrometry data analysis software. MS2extract also offers compatibility with GNPS2 MS/MS library batch submission. We also present PhenolicsDB (created with MS2extract), a phenolics-focused MS/MS library of 71 authentic standards with spectra collected in different polarities and various collision energies. This database is available for standalone download and use or can be accessed through GNPS2 for query and molecular networking applications. Finally, we use PhenolicsDB to improve the annotation of phenolic compounds in apple fruit as a use-case example of compound identification.

## Methods

In this section, we first describe the development and use of MS2extract. Then, we will explain how MS2extract was used to create an in-house high-resolution MS/MS library of analytical standards, PhenolicsDB, what it contains, and how it can be used. Finally, we describe a use-case of PhenolicsDB along with other publicly available GNPS2 MS/MS libraries for compound annotation in an untargeted metabolomics analysis of apple fruit.

### Software implementation

The impetus for the development of MS2extract was to provide an open-source tool to enable users to create in-house MS/MS libraries that could be used locally and/or shared with the community, or made available in repositories (e.g., GNPS2). MS2extract has four main steps (Figure 1): (a) importing the MS/MS data, (b) extraction of a specific precursor ion mass, (c) detection of associated MS/MS ions and removal of background noise, and (d) exporting the MS/MS library. This core pipeline is the backbone for the MS2extract “batch mode”, where iteration across different masses/files produces a multi-compound library. The batch mode functions can be distinguished from the core pipeline by the “batch_*” notation before the function’s name.

**Figure 1.**
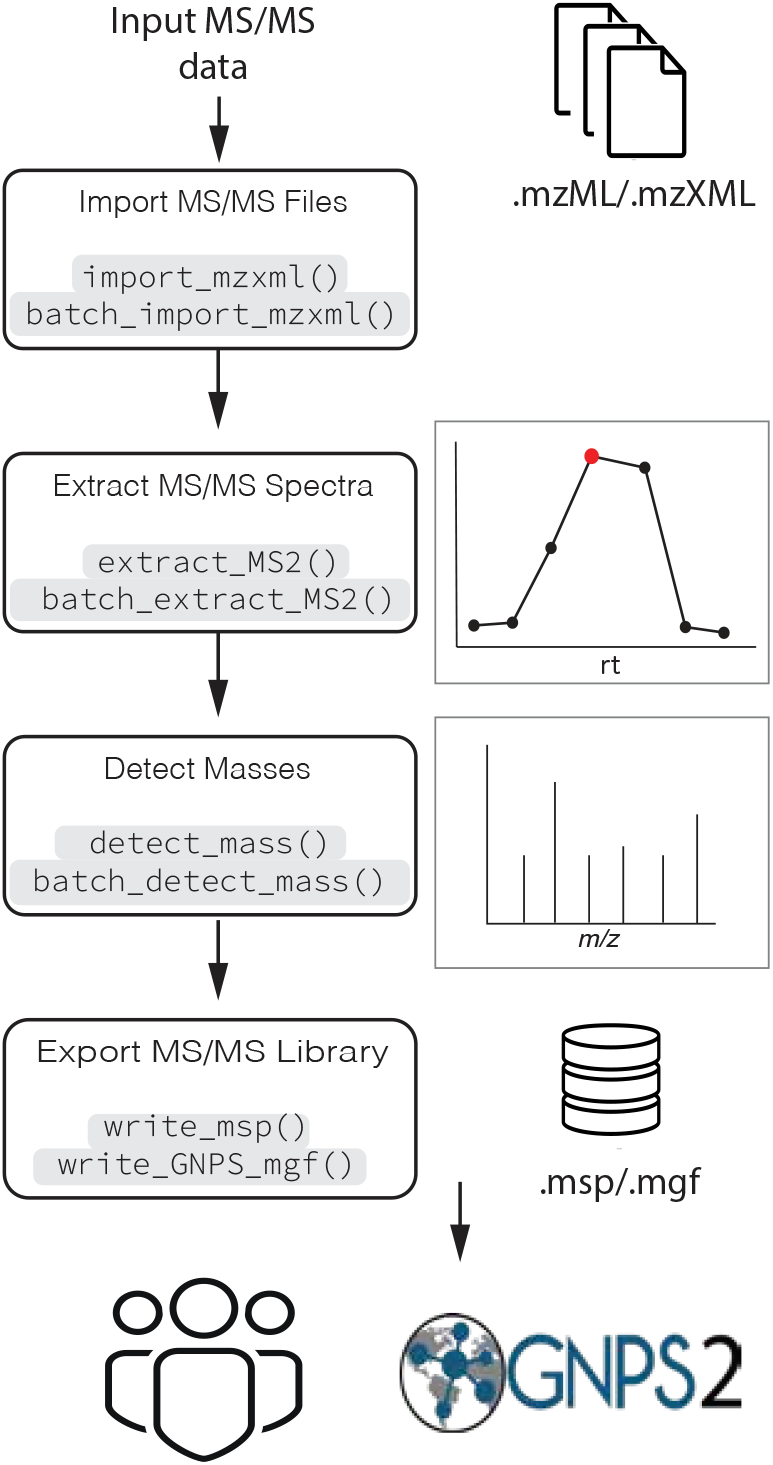
An overview of the MS2extract module for the LC-MS/MS library creation process. This process has four main steps: import MS/MS files, extract MS/MS spectra, detect masses, and export MS/MS libraries. R functions for single and batch functions are denoted highlighted in grey.

Additionally, MS2extract includes plotting functions to display helpful graphics (e.g., MS/MS total ion chromatograms (TIC) and spectra) throughout the processing pipeline. Supplementary Table 1 lists the MS2extract functions, their arguments, and descriptions of these arguments.

### Importing MS/MS data

MS/MS data can be imported using the import_mzxml() and batch_import_mzxml() functions. These functions require three arguments: *file*, the .mzML/.mzXML file path; *met*_*metadata*, a table with information about each metabolite, including compound name, chemical formula, and ionization mode; and *ppm*, a mass error tolerance for the precursor ion extraction window.

Importing raw MS/MS data has two parts: loading data in memory and filtering MS/MS scans. First, raw MS/MS data is loaded into memory using the MSnbase package^12^, then the data is formatted with the package masstools^13^, following tidy data principles^14^. The collision energy of each scan is then extracted with the mzR package^9^. Based on the metabolite information provided in *met_metadata*, the theoretical monoisotopic mass is calculated using the chemical formula with the Rdisop package^15^. Then, with the indicated ionization mode, the mass of a proton is added (positive polarity) or subtracted (negative polarity) from the monoisotopic mass to calculate the theoretical charged mass (*m/z)*. Finally, we used the equation for mass error (Eq. 1)^16^ to solve for *m*_*E*_, obtaining (Eq.2), where ppm is the error mass expressed in parts per million, *m*_*E*_ is the observed experimental mass (*m/z*), and *m*_*T*_ is the theoretical exact charged mass (*m/z*). With the resulting equation (Eq. 2), MS2extract calculates the mass error range (using symmetrical mass error, meaning 5 ppm will provide a range that is ±5 ppm) using a negative ppm value for the lower limit, and a positive value for the upper mass error limit.

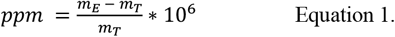

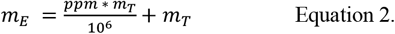

For example, 5*-O*-caffeoylquinic acid (i.e., chlorogenic acid), with a chemical formula of C_16_H_18_O_9_, in positive ionization polarity, has a *m*_*T*_ of 355.1023 *m/z*. Using a five ppm mass error tolerance, the lower mass limit is 355.1005 *m/z*, and the upper limit is 355.1041 *m/z*. With this calculated *m*/*z* range, scans with precursor ions falling within these values are kept (i.e., 355.1005 < precursor ion < 355.1041), while others are discarded. It is worth emphasizing that this calculation is based on a symmetric mass error tolerance, so the value given is both added and subtracted from the theoretical mass. This means that a value supplied of ± 5 ppm will lead to a 10 ppm window.

import_mzxml() also accepts a retention time window as *rt_min* and *rt_max* arguments to specify an elution window for each compound. This allows for the separate extraction of isobaric metabolites in the same datafile, given that they are resolved chromatographically.

### Extraction of representative MS/MS scans

MS2extract first extracts an MS/MS total ion chromatogram (TIC) for a given precursor mass. Then, it selects the most intense scan for downstream processing. The reason for this is that the most intense scan has the highest signal, which often results in the best signal-to-noise ratio. This same approach can be seen used in MZmine^11^.

By using the verbose argument (i.e., *verbose* = TRUE), users can decide whether the MS/MS TIC and spectrum are printed. In these plots, the mass of the precursor ion is indicated with a blue diamond if the precursor ion is present in the MS/MS spectrum, and with a white diamond if the precursor ion is absent. Additionally, extract_MS2() and batch_extract_MS2() automatically detect if one or multiple collision energies are present. When multiple collision energies are present, the most intense scan per collision energy is selected.

### Mass detection

The process of mass detection removes low-intensity product ions before exporting the MS/MS spectra to a reference library. In this step, users define a minimum signal intensity threshold, and if ions are below this threshold, they are removed. Mass detection is performed by functions detect_mass() and batch_detect_mass(). These functions take two arguments: *normalize* (accepts TRUE or FALSE), which governs whether a spectrum is normalized by dividing each product ion intensity by the base product ion peak (*normalize* = TRUE) or not (*normalize* = FALSE); and *min_int* (accepts a number), which indicates the minimum ion intensity threshold. By default, MS/MS spectra are normalized, and the minimum intensity cutoff is set to 1%. This default is based on Li et al.’s (2021)^17^, who found that removing intensities below 1% leads to more similar entropy scores between GNPS and MassBank for the same metabolite. Moreover, it is essential to emphasize that the normalization and minimum intensity cutoff values must be in concordance, since a normalized signal intensity can range from 0 to 100%, while a raw signal intensity can range from 0 to the maximum ion count observed.

### Export MS/MS libraries

Users can export libraries in .msp or .mgf formats by providing metadata about each small molecule, allowing appropriate metadata to be paired with the corresponding MS/MS peak list. To ensure a unique identifier for each library entry, an InChIKey is required. Both supported MS/MS library formats (.mgf and .msp) were checked and have compatibility with MZmine 4.0v, MS-DIAL 5.0v, SIRIUS 5.8v, and matchms 0.27v.

The .mgf format was aligned to be compatible with the GNPS batch upload of annotated spectra to enable a straightforward submission.

### Analytical standards and MS/MS data collection

#### Analytical standards and reagents

Seventy one analytical standards (organic and phenolic acids, dhydrochalcones, flavonoids aglycones, flavonoid glycosides, procyanidins, anthocyanins, and sugars) were purchased from Merck and Cayman Chemical (Supplementary Table 2). All standard solutions were prepared with H_2_O:MeOH in a 1:1 ratio and stored at -80 °C until data acquisition. Mass spectrometry grade acetonitrile and formic acid were acquired from Fisher Scientific. Mobile phases were ultra-pure (i.e., milliQ) water with 0.1% formic acid (A), and acetonitrile with 0.1% formic acid (B).

### Data Acquisition

Analytical standard solutions were analyzed on an ultra-high performance liquid chromatography system (Agilent (Santa Clara, CA, USA) 1290 Infinity II) coupled to a quadrupole time-of-flight mass analyzer (LC-QTOF, Agilent 6545 or 6546) with electrospray ionization (ESI). The column was a Waters (Milford, MA, USA) Acquity UHPLC HSS T3 column (2.1 x 50 mm, 1.8 μm particle size) kept at 40 °C. We used a chromatographic method previously developed by our group^18^. Briefly, the system was run at 0.5 mL per minute, and the gradient was as follows: 0-0.5 min 0% B; 0.5-8 min linear increase to 100% B; 8-9 min hold 100% B, 9.01-10.0 isocratic at 0% B, for a total of a 10 min run.

MS parameters were set as follows: gas temp 350 °C, gas flow 10 L/min, nebulizer 35 psig, sheath gas temp 375 °C, sheath gas flow 10 L/min, nebulizer 35 psig, sheath gas temp 11 L/min, VCap 4500 V, nozzle voltage 500 V, fragmentor1 100, skimmer1 45, octopoleRFPeak 750. Data was collected separately in both positive and negative ionization modes. Targeted MS/MS methods were created for each metabolite, where the theoretical product ion mass for each metabolite was calculated based on its chemical formula and adjusted for ionization polarity.

All analytical standards have MS/MS data at 20 and 40 eV collision energies, and some additionally have data at 60 and 80 eV.

### Data conversion

Raw MS/MS data was collected in .d format and was converted to .mzML format using MSconvert 3.0.2.2v.

The data was converted from profile to centroid mode, only scans at MS level 2 were kept, and each chromatogram was cropped to capture the retention time where each metabolite eluted.

Three hundred and twenty .mzML files were created and are available through the PhenolicsDB R data package. All the analytical standards in PhenolicsDB are reported in Supplemental Table 2, along with their observed retention times, InChIKey, and PubChem identifier. Information about ionization polarity and collision energies are also included.

### Case example: apple fruit untargeted metabolomics

We used pooled methanol extracts from a diversity of apple fruits^18^, with the goal of identifying as many phenolic compounds as possible from apple fruit. We collected data-dependent MS/MS data in positive and negative modes, at 20 and 40 eV. Then, raw MS/MS data in .d format was converted to .mzML, and peak processing and compound identification were conducted in MZmine 3.9v via the processing wizard pipeline with default parameters. Five GNPS2 MS/MS libraries (GNPS-library, MASSBANK, SUMNER, GNPS-natrual_products, GNPS-nutrimetabolomics), and PhenolicsDB were used for local compound annotation. Although PhenolicsDB includes retention times based on Bilbrey et al. (2021)^18^ chromatography method, we decided not to constrain feature annotations to match retention times, as we wanted to simulate a more typical scenario where retention time is unavailable, or data was collected using a different chromatographic method. Similarly, we decided to keep only the top annotation (i.e., identification with the highest score), for the downstream identification statistics summary. In cases where an annotation is present in both libraries (GNPS2-libraries and PhenolicsDB), the annotation is attributed to the library with the highest cosine score.

## Results and discussion

### MS2extract software

A comparison of the functionality of MS2extract currently available tools is presented in Table 1. There are three main advantages of using MS2extract: (a) batch functionalities for MS/MS library creation, (b) compatibility with GNPS2 library submission, and (c) pipeline customization. Batch functionalities are essential for processing large numbers of reference spectra at once, without needing to combine individual spectra later into one larger library. The only open-source software currently providing batch functionalities is MetID. However, its pipeline is designed as a one-stop solution, where the data is provided, and its pipeline carries out all processes, without any additional input. While the ease of use is high, it does not allow for further tailoring of the spectrum extraction and filtering process to meet specific needs. Specifically, users cannot set noise thresholds based on their instruments, which prevents the tailoring of the removal of low-intensity ions attributed to background noise. The presence of noise can artificially deflate MS/MS matching scores, making a library potentially less useful.

**Table 1.** Software benchmark for MS/MS library creation from raw data modules.

| Software | Dedicated MS/MS library creation module | Batch functionalities | Available GUI | Programming language | GNPS compatible | Pipeline customization | Ref |
| --- | --- | --- | --- | --- | --- | --- | --- |
| MS2extract | Yes | Yes | No | R | Yes | Yes | Present work |
| MZmine | Yes | No | Yes | Java | Partial | No | 11 |
| MetID | Yes | Yes | No | R | No | No | 10 |
| matchms | No | No | No | Python | No | Yes | 7 |
| CompoundDb | No | No | No | R | Yes | No | 19 |

On the other hand, GNPS2 has become one of the most popular MS/MS analysis environments, providing MS/MS-based feature annotation and other tools such as feature-based molecular networking, ion identity molecular networking, and spectrum resolver. Therefore, allowing easy compatibility for GNPS2 MS/MS library submission enables greater library reuse and spectral democratization. The only software providing GNPS2 compatibility is MZmine, though reference spectra cannot be created in a batch.

Furthermore, it is worth noting that MS2extract is a software dedicated to MS/MS library creation from raw MS/MS data, and it is not intended to be a general framework for mass spectrometry data handling and analysis, like matchms, openMS, or contain modules for handling already existing MS/MS libraries such as CompoundDb^19^. We chose R for the development of MS2extract to have a lower barrier for use, as R has a less steep learning curve when compared to other programming languages, and is already widely used by chemists and biologists.

### PhenolicsDB

Once MS2extract was developed, we used the batch processing mode to generate 320 reference MS/MS spectra described in Supplementary Table 2. We made our in-house MS/MS library publicly available in both .msp and .mgf formats. The library can be used through the PhenolicsDB R data repository (https://www.cooper-stonelab.com/PhenolicsDB/), embedded within GNPS2, or via download from a Zenodo repository^20^. PhenolicsDB has also been used in other projects, including to identify compounds in plants used in traditional medicine used in the Amazon region of Ecuador^21,22^, and from a mixed fruit and vegetable juice^23,24^.

Based on the uniqueness of InChiKeys present in PhenolicsDB and GNPS2 libraries, PhenolicsDB contributes with spectra of twelve unique compounds previously not present in GNPS2 (2,3-dihydroxybenzoic acid, 2,4,6-trihydroxybenzaldehyde, 8-hydroxykaempferol, 6-hydroxy-7-methoxycoumarin, procyanidins A2 and B3, procyanidin, quercetin 3-*O*-sophoroside, quercetin 3-*O-* glucoside, 4’-O-methyl quercetin, phloretin 4’-O-glucoside, and citramalic acid). We hope the addition of new spectra not previously present in GNPS2 may enhance metabolite identification in future studies.

### Case example: Application of PhenolicsDB to apple fruit untargeted metabolomics data

In untargeted metabolomics experiments, users ultimately want to identify the metabolites present in their samples. Knowing this, we aimed to develop a use case of the tool we report here, and referenced PhenolicsDB to see if it would help us identify more metabolites in our apple samples, over tools that are currently available. We collected data-dependent MS/MS data of methanolic apple fruit extracts and cross-referenced them against GNPS2 libraries (natural products focused) and PhenolicsDB. Using GNPS2 libraries and PhenolicsDB, we were able to annotate 89 features (Supplementary Table 3) (Figure 2). In the upset plot shown in Figure 2A, we present the intersection of annotations according to their polarity and collision energy. Most annotations, 73 annotations or 82%, were only found at a single polarity and collision energy (29, 28, 9 and 7 features were annotated in positive polarity at 40 and 20 eV, and negative polarity at 40 and 20 eV, respectively; Figure 2A), while 18% of annotations were found in at least two polarities or collision energies (16 annotations, Figure 2A). For example, in positive mode, 9 features were annotated at both collision energies, while only 1 feature was annotated in both polarities and at both collision energies.

**Figure 2.**
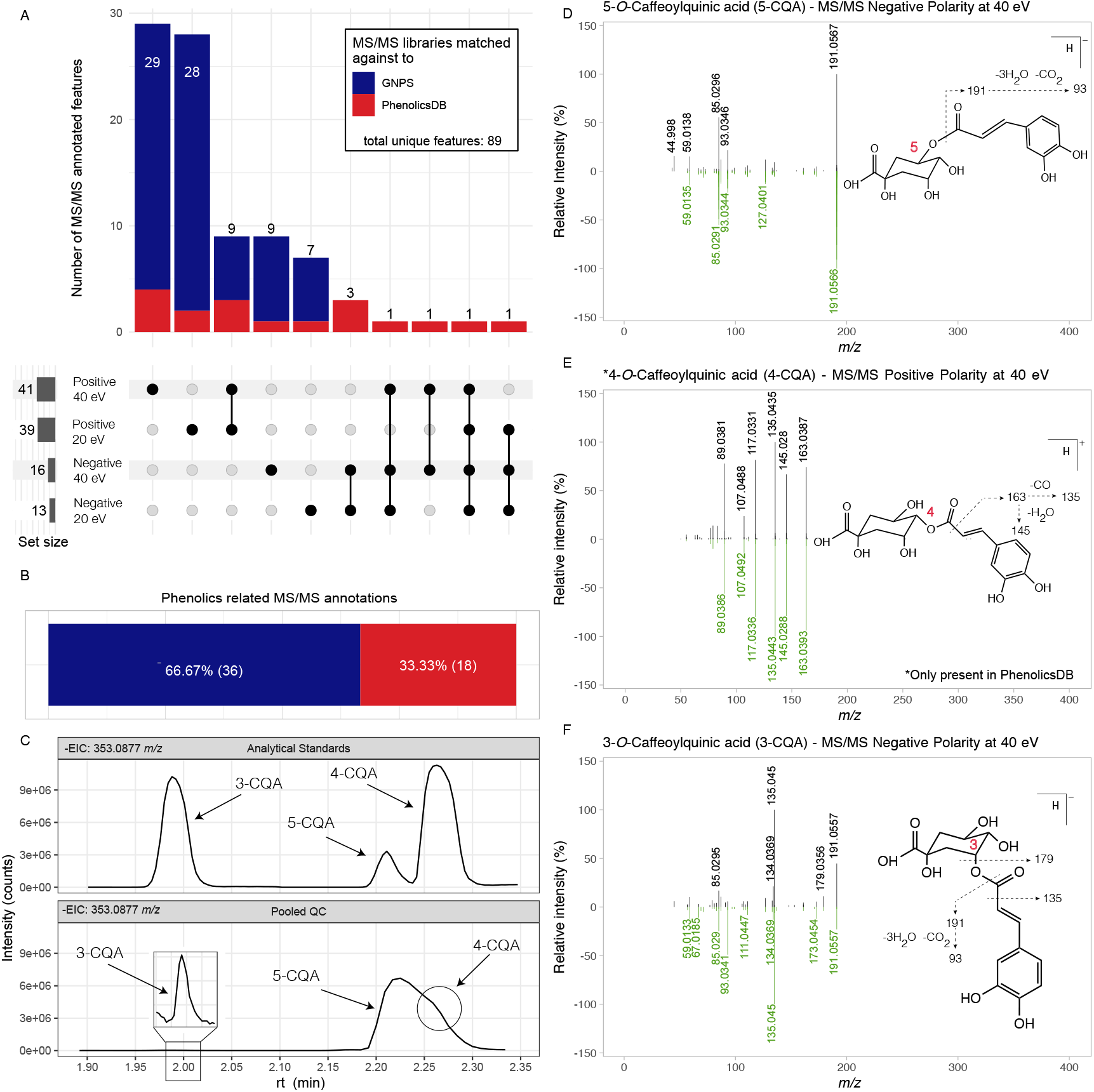
MS/MS annotations of apple fruit metabolites using GNPS2 MS/MS libraries and PhenolicsDB. (A) Upset plot for MS/MS annotations in negative and positive polarities in 20 and 40 eV, differentiated by the library to which each feature was matched against to. Phenolics-related MS/MS annotations. (B) Source of phenolic MS/MS annotations by library match. (C) EIC chromatogram for [M-H]^-^ of caffeoylquinic acid isomers between analytical standards (top) and pooled QC sample (bottom). Mirror plot for MS/MS annotation of 5-CQA (D), 4-CQA, which is only available in PhenolicsDB* (E), and 3-CQA (E) between the experimental spectrum and reference PhenolicsDB spectrum.

Although 74.2% (66) of unique features were annotated only in positive polarity (positive at 20 eV: 29, positive at 40 eV: 28, and positive at 20 and 40 eV, 9)(Figure 2A), the remaining features (23 features) were only annotated in negative mode. This finding was unexpected since most phenolic compounds are well documented to have better ionization efficiency in negative compared to positive mode. It is worth noting that the GNPS2 libraries used locally for compound annotation contain twice as many spectra in positive polarity (10,078 spectra) compared to negative polarity (4,986 spectra). We suggest that our higher number of annotated features in positive polarity resulted from more reference spectra in that mode.

Since PhenolicsDB is enriched with phenolic compounds, we expect its addition to improve the annotation of phenolic compounds. Out of the 89 identified features, 54 were phenolic annotations (Figure 2B). Among these 54 phenolic annotations, 36 (66.67%), were based on a GNPS2 match, while the remaining phenolic annotations (18 or 33.33%) were based on a PhenolicsDB match. This finding suggests that although phenolic compounds are present in MS/MS libraries, 20% more features (based on all 89 identified features), or 50% more features (based on 36 phenolic annotations) can be annotated when adding PhenolicsDB. Our findings are consistent with phenolic annotations that have been previously reported in apples^25,26^. We did not identify any phenolics (or compounds) in apple that had not been reported previously in literature^27,28^, though this is not surprising given its common study.

In the case of features matched against PhenolicsDB, we validated the annotations by referencing their observed retention time against retention time reported in PhenolicsDB (obtained from analytical standards) (Supp. Table 3). We found features annotated as apigenin (cosine score = 0.746 and rt = 2.90 min) and 4’-*O*-methylquercetin (cosine score = 0.775 and rt = 3.19 min) that did not have the same retention times as the authentic standards ( 3.94 and 4.08 min for apigenin and 4’-*O*-methylquercetin (Supp. Table 3), respectively). In both cases, the MS/MS spectra present characteristic flavonoid fragments, which suggest these two features can be structural isomers of the corresponding annotated phenolic. These two annotations, despite having cosine scores greater than 0.70, are false positives, giving a false discovery rate of 11% (2 out of 18 annotations). It is worth noting that this false discovery rate might be skewed to a lower rate as we collect MS/MS data for our samples and PhenolicsDB under the same conditions, and demonstrates the importance of ground truthing identifications.

For features annotated against GNPS2 libraries, which are also present in PhenolicsDB, we corroborated their retention time to validate their accurate annotation. We found six features (procyanidin B1, caffeic acid, quercetin 3-*O*-galactoside, phloretin, quercetin 3-*O*-rutinoside and, quercetin 3-*O*-rhamnoside) that were annotated against GNPS2 libraries, which are also present in PhenolicsDB (Supplementary Table 4.). From these six annotations, four annotations (caffeic acid, quercetin 3-*O*-galactoside, -3-*O*-rhamnoside, and -3-*O*-rutinoside) correctly matched with the retention time in PhenolicsDB, while two annotations (procyanidin B1 and phloretin) did not match their corresponding retention time. Therefore, we demonstrate that it is imperative to validate candidate annotations, when possible, for accurate compound reporting.

On the other hand, we cannot easily validate annotations matched against GNPS2 libraries only, without acquiring those authentic standards and running them with our methods, as they do not have retention time information, nor can we replicate data acquisition conditions due to the lack of chromatography method information. Moreover, after a closer inspection of GNPS2 annotations, we realized that a feature deemed as austinoneol (positive polarity at 40 eV, cosine score: 0.820), is likely a false positive, as austinoneol is only described to be synthesized by fungi^29^, and there is no literature available on its presence in plant sources.

It is worth highlighting that the representation of reference spectra in multiple acquisition conditions (i.e., polarity and collision energy) in MS/MS libraries could enable higher annotation rates. For example, of the caffeoylquinic acid (CQA) positional isomers: 3, 4, and 5-*O*-CQA, only the 3-CQA and 5-CQA isomers were found to be in the top candidate annotation spot when only using GNPS2 libraries. Moreover, after exploring all possible candidate annotations for all features with a cosine score greater than 0.70, a feature detected in positive polarity at 40 eV with the top 1 annotation as NCGC00180756 (cosine score = 0.809), was also annotated as 4-CQA as a fourth candidate annotation (cosine score = 0.806). However, when PhenolicsDB was incorporated as a reference library, 3- and 5-CQA isomers were still annotated as the top candidate annotations, and 4-CQA was annotated correctly as the top candidate with a cosine score of 0.915 (0.106 score units greater than the top annotation with GNPS2 libraries only), only in positive polarity at 40 eV (Figure 2E). Therefore, although features can be annotated against GNPS2 libraries, we improved the annotation rate with greater cosine scores and, with a low false positive rate, using PhenolicsDB. This improvement can be attributed to the availability of reference spectra under different acquisition conditions (polarity and collision energy), which PhenolicsDB offers.

The annotation of caffeoylquinic acid isomers becomes more accurate when retention time is incorporated (rt information available in PhenolicsDB), and the extracted ion chromatogram of the caffeoylquinic acid positional isomers mass in negative polarity (*m/z* 353.0887) of the pooled apple quality control sample is displayed against the authentic analytical standards (Figure 2C). While the 3-CQA peak is fully resolved at 1.99 min, the 4- and 5-CQA isomers closely elute with retention times of 2.26 and 2.21 min, respectively. However, due to the combination of acquisition conditions, and the 5-CQA isomer being the predominant chemical species in the apple fruit, with peak broadening, the resolution between these two isomers (5-CQA and 4-CQA) is decreased.

The MS/MS mirror plots for the annotation of 5-, 4-, and 3-CQA positional isomers are presented in Figure 2D, 2E, 3F, respectively. It is worth noting that due to the 4-CQA and 5-CQA coelution, the experimental spectrum of 4-CQA (Figure 2F), also contains the contribution of 5-CQA product ions. However, this MS/MS annotation was only possible as the DDA engine triggered an MS/MS event (only recorded in positive polarity at 40 eV) at a retention time that also included 4-CQA product ions. The identification of these three caffeoylquinic acid isomers (3-, 4-, and 5-CQA) is crucial for improving metabolome coverage, as the 3- and 4-CQA isomers are often not reported or studied due to the low concentration in apple^30,31^, and one of the reasons might be the 5-CQA isomer signal might obscure the signal of the other isomers.

This case example showcases the benefit of improving the metabolome coverage and the identification of the apple metabolome aided by public MS/MS libraries and PhenolicsDB. Additionally, with the available retention time information in PhenolicsDB, researchers can utilize the reported retention times of authentic standards, if chromatography conditions are the same, to enhance the level of identification from level 2 (MS/MS match) to level 1 (MS/MS and retention time match).With these two sources of orthogonal information, we hope researchers who use our PhenolicsDB can increase the confidence in compound identification and reporting.

## Conclusions

Our proposed software was suitable for creating and automating reference MS/MS libraries. Our framework imports vendor-independent data formats (.mzML and .mzXML), making it appropriate for a more comprehensive set of applications. However, further testing will be necessary to ensure compatibility with different mass analyzers. By making MS2extract able to export MS/MS libraries in open-source formats (.mpg and .mgf), and its compatibility with GNPS2 batch submission, we hope we can incentivize MS/MS library democratization. Moreover, using MS2extract batch functions, we successfully created PhenolicsDB, allowing us to process more than 320 reference MS/MS spectra of analytical standards. Subsequently, PhenolicsDB was made publicly available for download and use in multiple repositories, showing the potential of our framework to be compatible with GNPS2 MS/MS library submission and maximizing the potential use in every single GNPS2 task for feature annotation. PhenolicsDB also contains retention time information for each analyte that could be used to reach a higher confidence level of analyte identification, as previously discussed for the 3-, 4-, and 5-CQA positional isomers.

## ASSOCIATED CONTENT

### Supporting Information

Supplemental tables 1-4 are available on the publisher’s site.

MS2extract: https://www.cooperstonelab.com/MS2extract/

PhenolicsDB: www.cooperstonelab.com/PhenolicsDB/

## AUTHOR INFORMATION

### Author Contributions

CDQM and JLC co-developed the concept, CDQM acquired MS/MS data, CDQM developed the R package with input from JLC, CDQM and JLC developed documentation materials, CDQM drafted the manuscript, JLC edited the manuscript, and JLC has responsibility for the final manuscript content. All authors have approved the final version of the manuscript.

### Notes

The authors have no competing financial interests to declare.

## ACKNOWLEDGMENT

This work was supported by USDA AFRI (2023-67013-40267) and Hatch funds (OHO01563), and the Foods for Health Initiative, a focus area of the Discovery Themes at Ohio State University.

